# Atrial t-tubules adopt a specialist developmental state while alterations to Ca^2+^ buffering maintain systolic Ca^2+^ during postnatal development

**DOI:** 10.1101/2023.10.01.560329

**Authors:** CER Smith, JD Clarke, CJ Quinn, Z Sultan, H Najem, NC Denham, DC Hutchings, GWP Madders, JL Caldwell, LK Toms, C Pinali, DA Eisner, AW Trafford, KM Dibb

**Affiliations:** Division of Cardiovascular Sciences, Unit of Cardiac Physiology, School of Medical Sciences, Faculty of Biology, Medicine and Health, The University of Manchester, Manchester Academic Health Science Centre, 3.14 Core Technology Facility, 46 Grafton Street, Manchester, M13 9NT, United Kingdom

## Abstract

Transverse (t)-tubules ensure a uniform rise in calcium (Ca^2+^) and thus contraction in cardiac cells. Though more extensively studied in the ventricle, t-tubules also play a key role in the atria of large mammals, such as human, and their loss in heart failure is associated with impaired Ca^2+^ release and thus contractility. T-tubule restoration is therefore an ideal therapeutic target but the process of t-tubule formation is not understood. The aim of this study was to determine how t-tubules develop in the healthy atria and the impact this has on Ca^2+^ handling. Postnatal development was assessed in sheep from newborn through to adulthood. Atrial t-tubules were present at birth in the sheep atria and increased in density up until 3 months of age. In the latter part of development (3 months to adult) a lack of t-tubule growth but increase in cell width results in t-tubule density decreasing. In the newborn, despite reduced t-tubule density, we found the amplitude of the Ca^2+^ transient was maintained and this was associated with increases in the L-type Ca^2+^ current (*I*_Ca-L_) and the Ca^2+^ content of the sarcoplasmic reticulum (SR). We suggest these changes are sufficient to overcome the elevated cytosolic Ca^2+^ buffering in the newborn and the decreased t-tubule density. We have shown the neonate atria is highly specialised to negate reduced central Ca^2+^ release through enhanced surface *I*_Ca-L_ and SR load. This maintains atrial function despite immature t-tubules highlighting important differences in Ca^2+^ handling in the newborn and heart failure atria where t-tubules are sparse.

## 1. Introduction

Transverse (t)-tubules play a key role in facilitating a uniform rise in calcium (Ca^2+^) throughout the cell. They contain L-type Ca^2+^ channels (LTCC) that closely couple to ryanodine receptors (RyR) of the sarcoplasmic reticulum (SR) ensuring synchronous Ca^2+^ release throughout the myocyte. T-tubules are found in all mammalian ventricular myocytes[1, 2] and in the atria of large mammals[3–7] where they play an important role in triggering Ca^2+^ release similar to the ventricle[5, 8, 9]. Heart failure (HF) results in extensive t-tubule remodelling, particularly in the atria[2, 5]. Similarly to some studies in the ventricle[10–12], we have previously shown that atrial t-tubule loss in HF is associated with reduced LTCC current (*I*_Ca-L_) and consequently reduced systolic Ca^2+^ release[13]. As the contribution of atrial contraction to ventricular filling increases in HF[14], atrial t-tubule loss likely contributes to overall cardiac dysfunction. As such, the restoration or preservation of t-tubules, including those in the atria, would be beneficial therapeutically. However, despite the importance of t-tubules in triggering central Ca^2+^ release in the atria of large mammals, little is currently known about the process by which atrial t-tubules develop.

Ventricular t-tubules are absent or sparse at birth and develop postnatally resulting in changes in excitation-contraction (EC) coupling [8, 9, 15–21]. In the rabbit ventricle the absence of t-tubules in early life means LTCCs occur exclusively on the surface sarcolemma[22] resulting in reduced *I*_Ca-L_, impaired coupling with the SR and thus decreased systolic Ca^2+^ release[15, 16, 23–26]. In the newborn rabbit ventricle Ca^2+^ influx via reverse-mode operation of the sodium-calcium exchanger (NCX) is sufficient to generate contraction, compensating for reduced *I*_Ca-L_ and bypassing SR-mediated Ca^2+^ release[27–30]. As t-tubules develop, *I*_Ca-L_ and the reliance on SR Ca^2+^ release increases bringing about a more uniform rise in Ca^2+^ and a gain in EC coupling[15, 25, 31].

In newborn atrial cells we have shown enhanced Ca^2+^ release sites occur on the cell surface which we suggest helps to maintain the systolic Ca^2+^ transient when t-tubules are sparse [9]. While ventricular t-tubule development has been described, how atrial t-tubules develop postnatally and their influence on atrial EC coupling is unknown. By examining how t-tubules develop and the concomitant changes in Ca^2+^ handling in the atria, we sought to understand the factors that maintain the systolic Ca^2+^ transient when the t-tubule network is sparse in the newborn and extensive in the adult.

Here we show the time course for t-tubule development and Ca^2+^ handling changes in the atria. We report that in early life, when t-tubules are sparse, the Ca^2+^ transient amplitude is maintained by elevated *I*_Ca-L_ and SR load that offset increased intracellular Ca^2+^ buffering. As t-tubules develop they adopt a specialist developmental state, at 3 months post birth, characterised by increased t-tubule size and density which appear more complex in arrangement. Our data suggests increased t-tubules and triggered Ca^2+^ release allows *I*_Ca-L_ and SR load to decrease to adult levels; throughout this transition atrial cells are able to maintain the amplitude of the Ca^2+^ transient due to a decrease in intracellular Ca^2+^ buffering.

## 2. Methods

All procedures were carried out in accordance with The United Kingdom (Scientific Procedures) Act 1986, the European Union directive EU/2010/63 and the ARRIVE guidelines for reporting the use of animals in scientific experiments.

### 2.1 Sheep model of development

Sheep aged between 1 day and 1 week (newborn; NB), ∼1 month (1M), ∼3 months (3M), ∼6 months (6M) and adult ∼18 months of age (AD) were selected for inclusion in the study. Both males and females from a range of breeds were included. To track gross postnatal developmental changes, body weight and heart weight were recorded and a heart weight to body weight ratio calculated. In vivo cardiac function was assessed in conscious, non-sedated sheep using electrocardiography and echocardiography. Electrocardiographic measurements were taken using five surface electrodes and recorded for 10 minutes using IOX software (EMKA technologies). ECG recordings were converted from MKT to TXT files in ECG Auto (EMKA technologies) then analysed in Labchart (ADInstruments). Transthoracic echocardiographic images were obtained using a Sonosite Micromaxx (BCF Technology) or Vivid 7 (GE Healthcare) with short-axis 2D views at the mid-papillary muscle level recorded[32]. Images were converted using either VirtualDub or MicroDicom then opened in ImageJ (National Institutes of Health) for analysis. The lumen of the ventricle in diastole and systole in numerous cardiac cycles were drawn around using the polygon area tool and the left ventricular end diastolic area (EDA) and end systolic area (ESA) used to calculate fractional area change during contraction as a measure of contractility. Septal wall thickness (SWT) and posterior wall thickness (PWT) were also measured in diastole (d) and systole (s) on the same images used for EDA and ESA (Figure 2.1). Relative wall thickness was calculated for both SWT and PWT as a % change in thickness between diastole and diastole.

### 2.2 Isolation of atrial myocytes

Single myocytes were isolated from the left atrial appendage as described previously[2, 5, 13, 33]. Briefly, heparin (10,000 units) was administered to prevent clotting and animals were euthanased by pentobarbitone overdose (200 mg/kg). Hearts were removed, washed in Ca^2+^-free solution (containing in mM: 134 NaCl, 11 glucose, 10 HEPES, 10 2,3-butanedione monoxime, 4 KCl, 1.2 MgSO_4_, 1.2 NaH_2_PO_4_ and 0.5 mg/ml bovine serum albumin, pH 7.34 with NaOH) and the atria separated from the ventricles below the coronary sulcus. The left atrium was cannulated via the circumflex artery or a smaller branch of the circumflex if present and perfused at 37°C with Ca^2+^-free solution. Tissue was enzymatically digested using 0.3 mg/ml collagenase (Type II, Worthington or Type A, Roche) and 1.25 mg/ml pancreatin from porcine pancreas (Sigma) for developmental time points and 0.45-0.55 mg/ml collagenase and 0.05 mg/ml protease (Type XIV, Sigma) for adult animals. Following digestion, tissue was perfused then chopped and gently triturated in a taurine solution (containing in mM: 113 NaCl, 50 taurine, 11 glucose, 10 HEPES, 10 2,3-butanedione monoxime, 4 KCl, 1.2 MgSO_4_, 1.2 NaH_2_PO_4_, 0.1 CaCl_2_ and 0.5 mg/ml BSA, pH 7.34 with NaOH). Cells were filtered using a 200 µm nylon CellMicroSieve (Fisher Scientific) and resuspended in taurine solution prior to the reintroduction of Ca^2+^ by mixing taurine solution 50:50 with Tyrode’s solution (containing in mM: 140 NaCl, 10 HEPES, 10 glucose, 4 KCl, 2 probenecid, 1.8 CaCl_2_ and 1 MgCl_2_, pH 7.34 with NaOH). Cells were stored in Tyrode’s solution at room temperature prior to experimentation.

### 2.3 Confocal t-tubule imaging

Isolated atrial myocytes were stained with 4 µM di-4-ANEPPS for 10 minutes (Invitrogen) for membrane visualisation[3, 5]. T-tubule stacks were acquired using a Leica SP2 confocal microscope (488 nm excitation, 505 nm long pass emission filter) as described previously with an XY pixel size of 100 nm and Z-step depth of 162 nm[2, 3, 5, 11]. Images were deconvolved using Huygens Software (Scientific Volume Imaging) and a measured point spread function (PSF). Following deconvolution, images were thresholded to remove background fluorescence and a central sub-stack devoid of the top and bottom of the cell selected. The fractional area taken up by t-tubules within a cell as a measure of t-tubule density was calculated using a 2 µm projection binary projection in ImageJ where the number of black pixels (t-tubules) was expressed as a fraction of the total pixels within the cell[2, 11]. The distance 50% of voxels within the cell are to the nearest membrane (t-tubule or surface, t-tubule only and surface only) was also assessed using a custom IDL routine (Excelis VIS) and termed half-distance as described in detail previously[2, 3, 5, 11]. Here central image stacks were interpolated using the ‘congrid’ routine to give the same spacing in x, y and z and then the ‘morph_distance’ routine ran to create a distance map of voxel distance to membrane. To assess distance to t-tubules only and surface only the cell surface sarcolemma or t-tubules were removed respectively in ImageJ prior to running through IDL.

### 2.4 Electrophysiological recordings of intracellular Ca^2+^ and *I*_Ca-L_

To permit measurement of free intracellular Ca^2+^, [Ca^2+^]_i_, myocytes were loaded with 5 µM Fluo-5F AM (ThermoFisher) for 10 minutes and allowed to de-esterify for at least 30 minutes before experimentation. Measurements of [Ca^2+^]_i_ and membrane currents were made simultaneously under voltage clamp control using the perforated patch method with 240 μg/ml amphotericin-B added to a K^+^-based pipette solution (containing in mM: 125 KCH_3_O_3_S, 20 KCl, 10 NaCl, 10 HEPES, 5 MgCl_2_, pH 7.2 with KOH) [13, 35–37]. Cells were stimulated at 0.5 Hz using a 100 ms step from -40 to +10 mV and superfused with an experimental solution (containing in mM: 140 NaCl, 10 HEPES, 10 glucose, 5 4-AP, 4 KCl, 2 probenecid, 1.8 CaCl_2_, 1 MgCl_2_, 0.4 DIDS and 0.1 BaCl_2_, pH to 7.34 with NaOH). Probenecid was included to decrease indicator loss from the cell. A hyperpolarising step from -40 to -50 mV was applied to measure cell capacitance and thus surface area [38]. To assess SR Ca^2+^ content and buffering capacity, 10 mM caffeine was applied with the resultant Na^+^-Ca^2+^ exchange (NCX) current integrated and assessed alongside the caffeine-evoked Ca^2+^ transients as described previously [13, 33, 35–37, 39, 40]. The contribution of pathways other than *I*_NCX_ to Ca^2+^ removal during caffeine application was unaltered during development (Supplemental Figure 3). Therefore, we used our previously published NCX correction factor of 1.14 throughout this study[33]. Ca^2+^ buffering was assessed as previously described [33, 41], with the relationship between and total intracellular Ca^2+^ measured from the corrected integral of *I*_NCX_ and intracellular free Ca^2+^ obtained from Fluo-5F and fit with the following hyperbolic equation (equation 1):

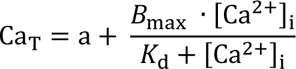

For which Ca_T_ is total Ca^2+^, *a* is an offset because the relationship cannot be determined for values of [Ca^2+^]_i_ below the resting level, *B*_max_ is the maximum buffering capacity for Ca^2+^, [Ca^2+^]_i_ is the free intracellular Ca^2+^ and *K*_d_ is the dissociation constant of the buffers. Ca^2+^ buffer power was calculated as previously described[33, 40] from the *B*_max_ and *K*_d_ values obtained above using equation 2:

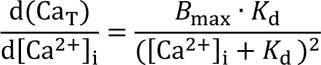

The increase of Ca_T_ during the Ca^2+^ transient (Δ[Ca]_Total_) was calculated from the resting and peak systolic [Ca^2+^]_i_ and the buffer properties.

### 2.5 Cell volume and nucleation

Isolated atrial myocytes were loaded with 20 µmol/l calcein-AM for 20 minutes at room temperature and image stacks acquired as described previously at an *xyz* resolution of 210 nm[13, 33, 35] (Supplemental Figure 2). Cell stacks were thresholded using half the maximum intensity of the plot profile and cell volume calculated from the number of voxels within the cell[35]. Surface area to volume ratios were calculated using a similar approach to [13, 33, 38] though as values could not be obtained from within the same cells the mean cell volume and cell capacitance for each time point was used (Supplemental Figure 2E). As calcein loading permitted visualisation of nuclei, the number of nuclei per cell was counted to assess the proportion of cell binculeation as a measure of developmental maturity.

### 2.6 Electron microscopy tissue processing and imaging

Left atrial appendage tissue was collected from newborn, 3 months and adult sheep and fixed in 2.5% glutaraldehyde and 2% paraformaldehyde in 100 mM sodium cacodylate buffer, pH 7.2. Following washing with sodium cacodylate buffer, samples were stained in 2% osmium tetroxide and 1.5% potassium ferrocyanide; 1% thiocarbohydrazide; 2% osmium tetroxide, 1% uranyl acetate and Walton’s lead nitrate with water washes between each step as previously described but with small adaptations [42–44]. Following staining, an ascending series of ethanol (50%, 70%, 90%, 100%, 100%) then propylene oxide was used to dehydrate samples prior to infiltration treatment with increasing concentrations of TAAB 812 hard resin mixed with propylene oxide (25%, 50%, 75%, 100%). Samples were then embedded in pure resin inside plastic moulds and baked for 36 hours at 60°C. For imaging, samples were cut from moulds, mounted onto cryo pins and coated in a gold-palladium alloy. Serial images were obtained at 50 nm intervals with a pixel size of 6-12.5 nm using a Quanta 250 FEG scanning electron microscope and Gatan 3View device operated at 3.8 kV and approximately 0.46 Torr. Image stacks were opened and t-tubule reconstruction and volumetric analysis was performed in 3DMod (IMOD) [43, 45] .

### 2.7 Immunoblotting

Left atrial appendage tissue was chopped into 1 mm^3^ chunks and rapidly snap frozen in liquid nitrogen. Tissue was homogenised in RIPA buffer containing protease and phosphatase inhibitors (0.1 mg/ml phenylmethanesulphonylfluoride, 100 mmol/l sodium orthovanadate, 1mg/ml aprotonin, 1mg/ml leupeptin) and protein concentration assessed by using a Bradford assay. Samples were run on NuPAGE 4-12% Bis-Tris gels (ThermoFisher) and then transferred onto nitrocellulose membranes (GE Healthcare) as described previously[2, 11, 13, 33]. Total protein was stained using Ponceau S (Applichem Lifescience) and membranes imaged prior to blocking in 5% BSA, SEABLOCK® (ThermoFisher) or 5% milk. Membranes were incubated with primary antibodies targeted to AMPII (1:200, Santa Cruz Biotechnology), JPH2 (1:5000, Santa Cruz Biotechnology) or TCAP (1:1000, Abcam) followed by HRP-conjugated anti-mouse, anti-goat and anti-rabbit secondary antibodies (1:40,000, Santa Cruz Biotechnology). Chemiluminescence substrate (GE Healthcare) was added for protein detection via digital capture (Genesnap, Syngene). All blots were run in triplicate, with target protein expression normalised to total protein levels and an internal control standard that was loaded on each gel as previously described[2, 10].

### 2.8 Statistics

Data was tested for normality using the Kolmogorov-Smirnov or the Shapiro-Wilk test (GraphPad Prism or SPSS statistics, IBM) and transformed if required. Statistical significance was assessed using one way ANOVA (GraphPad Prism), Chi-square (SigmaPlot), two way ANOVA or linear mixed modelling (SPSS Statistics, IBM) as appropriate, with mixed modelling used to account for multiple observations from the same animal. Data are presented as mean ± SEM for *N* animals and *n* cells or repeats, with these observations shown behind mean data on graphs as filled or open circles respectively. Statistical significance was attained when *p*<0.05, with * showing significant difference *vs* adult and $ showing significant difference *vs* previous time point.

## 3. Results

### 3.1 Atrial t-tubules grow until 3 months of age despite increasing cell width into adulthood

We firstly mapped development of the heart using echocardiography and the electrocardiogram (ECG; Supplemental Figure 1A). Left ventricular systolic function was maintained throughout development despite the change in heart size over the first 3 months (Supplemental Figure 1A). Heart rate decreased to 6 months of age while atrial electrical parameters, P wave duration and PR interval, increased to 3 months of age and then plateaued (Supplemental Figure1B). In terms of the ventricle, QRS duration did increase between newborn and adult but the corrected QT interval was unaltered with age (Supplemental Figure 1B). Taken together this data suggests maturation of the ventricle may occur before that of the atria.

As we have shown in newborn tissue (<1 day old; [9]) we found sporadic t-tubules present in isolated atrial myocytes from newborn animals up to 1 week of age (Figure1A). Both t-tubule density and cell width increased during development and as cells matured binucleation increased (Figure1A and Supplemental Figure 2A-C). The role of t-tubules is to minimise the distance between the cell membrane and the Ca^2+^ release machinery. Therefore we assessed t-tubule density by calculating the average distance of all points within the cell to the nearest membrane in the x, y or z dimension using distance maps as described previously (half distance; [2, 3, 5, 11]). The half distance to any membrane (t-tubule and surface; Figure 1Bi inset) decreased between newborn and 3 months of age indicative of t-tubule growth (Figure 1Bi; *p*<0.001). However, this reduction was reversed beyond 3 months into adult with surprisingly no difference in half distance to membrane observed between newborn and adult.

**Figure 1.**
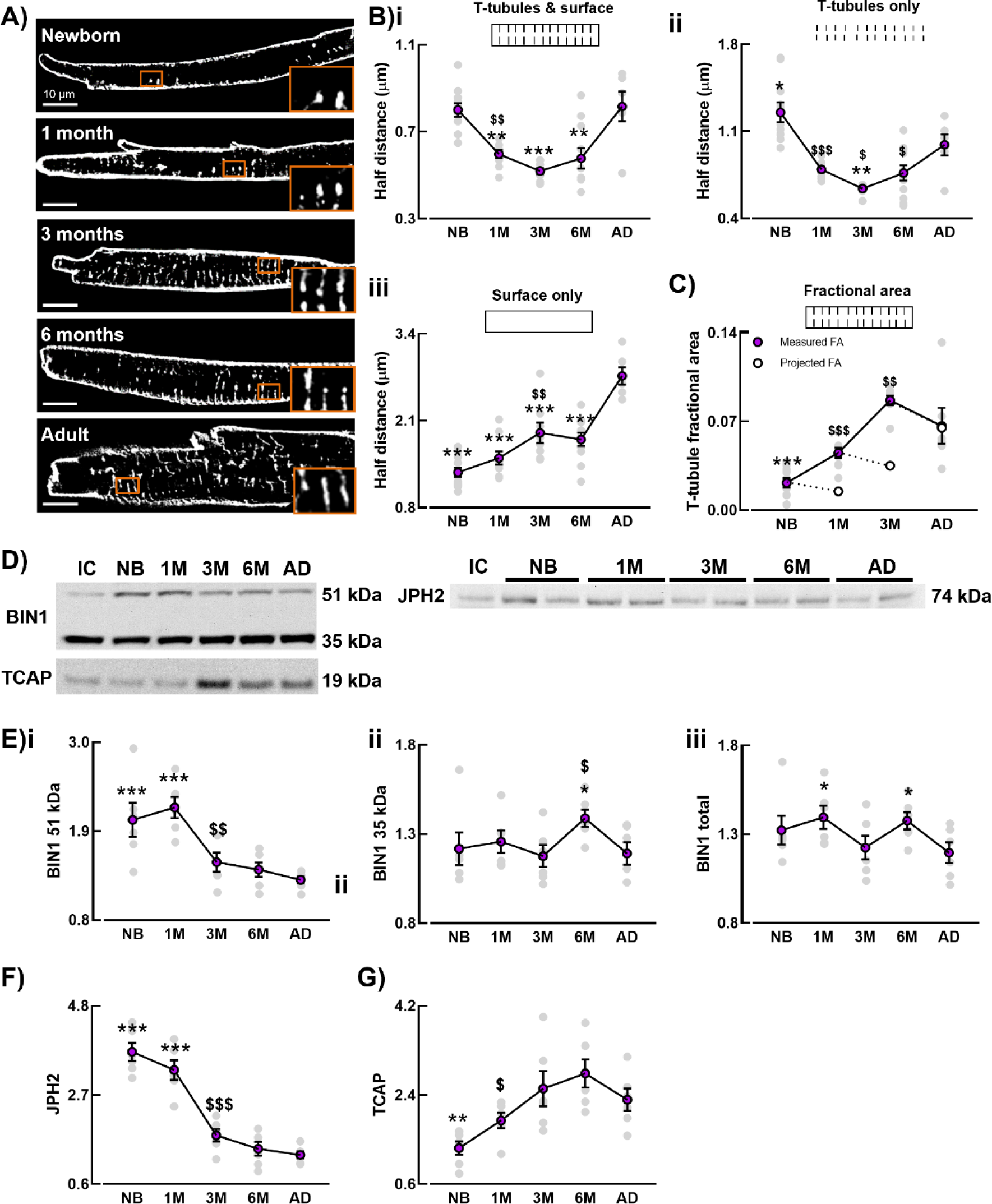
Developmental changes in atrial t-tubule density and expression of t-tubule regulatory proteins. Representative images of t-tubule staining in atrial myocytes (A). Mean data for half distance to membrane (t-tubules and surface) (Bi), half distance to t-tubule membrane only (ii) and half distance to surface membrane only (iii). Mean and projected fractional area should cell size increase but no further t-tubule formation occurs between time points (C). Representative blots (D) and mean data for expression of BIN1 (E), JPH2 (F) and TCAP (G). B&C, n=37-58 cells/N=6-11 animals. E-G, N=6 animals, minimum 3 repeats. * *p*<0.05, ** *p*<0.01, *** *p*<0.001 *vs* adult, $ *p*<0.05, $$ *p*<0.01, $$$ *p*<0.001 *vs* previous time point, using linear mixed models analysis. Scale bars = 10 µm.

We next separated the contribution of t-tubules and surface membrane to the half distance measurement (Figure 1Bii&iii). Consistent with increasing cell width throughout development the half distance to the surface membrane increased into adulthood (Figure 1Biii, *p*<0.001). Similarly to the half distance to all membranes, half distance to t-tubules also displayed a U-shaped relationship with age, suggesting t-tubule growth over the first 3 months predominated over the increase in cell width (Figure 1Bii, *p*<0.001). However, after 3 months of age the increase in half distance to t-tubules suggests increasing cell width predominates (Figure 1Bii). As expected, half distance is decreased in the adult *vs* newborn consistent with the increased importance of t-tubules. Thus, atrial t-tubule development is not directly related to increasing cell width. We next investigated if t-tubule growth stopped at 3 months of age despite cells still being immature.

The fractional area of the cell occupied by t-tubules increased to 3 months of age as t-tubules grew (purple symbols, Figure 1C). The white symbols show projected fractional area values assuming no t-tubule growth from the previous time point. For example, if the t-tubules present in newborn cells were put into a 1 month cell we would predict fractional area to decrease from 0.022 to 0.015. Instead, measured fractional area at 1 month was 0.045 thus showing continued t-tubule formation. At 1 and 3 months t-tubule fractional area was greater than predicted (purple>white) confirming t-tubule growth. The fact that the symbols overlay in the adult (purple symbol is behind the white symbol) suggests between 3 months and adult cell growth occurs without t-tubule growth. Therefore t-tubule growth stops at 3 months and the continued increase in cell size results in a decrease in t-tubule density between 3 months and adulthood (Figure 1C). This is consistent with the lack of change in cell capacitance between 3 months and adulthood despite increases in cell volume and width (Supplemental Figure 2C-D). To determine if t-tubule growth correlates with gross cardiac development we measured developmental changes in body and heart weight (Supplemental Figure 1C). Both parameters increased after t-tubule growth stopped (3 months) however the ratio between the two was unchanged after 3 months of age showing further growth of the heart and body occurred at the same relative rate.

### 3.2 BIN1, JPH2 and TCAP reveal distinct expression changes during t-tubule development

To understand if known t-tubule associated proteins could be involved in t-tubule growth we quantified the expression levels of BIN1 (also known as Amphiphysin-2), Junctophilin-2 (JPH2) and Telethonin (TCAP; Figure 1D-G). Two bands at 51 and 35 kDa were identified for BIN1 as previously reported[2]. Expression of both 51 kDa BIN1 and JPH2 were increased during the t-tubule growth phase but had decreased to adult levels once t-tubule were formed (3 months; Figure 1E&F; *p*<0.001). Changes in the 35 kDa band or total BIN1 did not correlate with t-tubule development (Figure 1Eii-iii). Unlike BIN1 (51 kDa) and JPH2, TCAP expression increased during early life (Figure G; *p*<0.01). Therefore, although based on an association, our data suggests a possible role for BIN1 (51 kDa isoform) and JPH2 in t-tubule formation or organisation in the dyad and TCAP in t-tubule extension and maintenance in the adult. Next, we examined the ultrastructure of developing atrial t-tubules to identify structural changes which underpin the increase in t-tubule density at 3 months.

### 3.3 Distinct t-tubule morphologies during development account for changes in t-tubule density and structure

Since confocal microscopy lacks the resolution to understand the detailed structure of t-tubules we used serial block face Scanning Electron Microscopy to resolve detailed atrial t-tubule ultrastructure (sbfSEM; Figure 2). T-tubules were manually reconstructed from sbfSEM images of newborn, 3 months and adult cells and typical examples are shown in transverse and longitudinal orientation (Figure 3A; t-tubules white and cell surface purple). T-tubules in newborn were usually sporadic and shorter in length compared to those in later life, however we did observe a single cell where t-tubules appeared almost fully developed at birth. Interestingly, the improved resolution revealed that t-tubules present at 3 months were more complex in structure compared to newborn and adult (Figure 2A). Similarly to confocal data, t-tubule density measured from sbfSEM images (t-tubule volume/10 sarcomeres) increased from birth to 3 months and declined to adulthood (Figure2B, *p*<0.001). Increasing t-tubule density occurred via an increase in both the number and volume of t-tubules while increased volume was due to increases in both t-tubule diameter and branching (Figure 2C-F; *p*<0.001). The decrease in t-tubule density into adulthood was not due to a change in t-tubule number (Figure 2C; *p*=0.99) but was associated with a non-significant decrease in t-tubule volume (Figure 2D; *p*=0.2) suggesting changes in t-tubule size or morphology were likely responsible. Consistent with this concept, decreases in t-tubule diameter (Figure 3E; *p*<0.05) and branching (non-significant, Figure 3F) in the adult suggests t-tubule dilatation or increased complexity at 3 months underlies the increased t-tubule density compared to the adult. Interestingly, t-tubule volume, diameter and branching returned to newborn levels in the adult suggesting t-tubule morphology at 3 months is distinct. To determine if this was the case, t-tubule morphology was categorised into mature structures (based on the most common morphologies seen in adult), t-tubule stumps (immature short t-tubule extensions) and disordered t-tubules which encompass complex, branched and more longitudinally orientated tubules.

**Figure 2.**
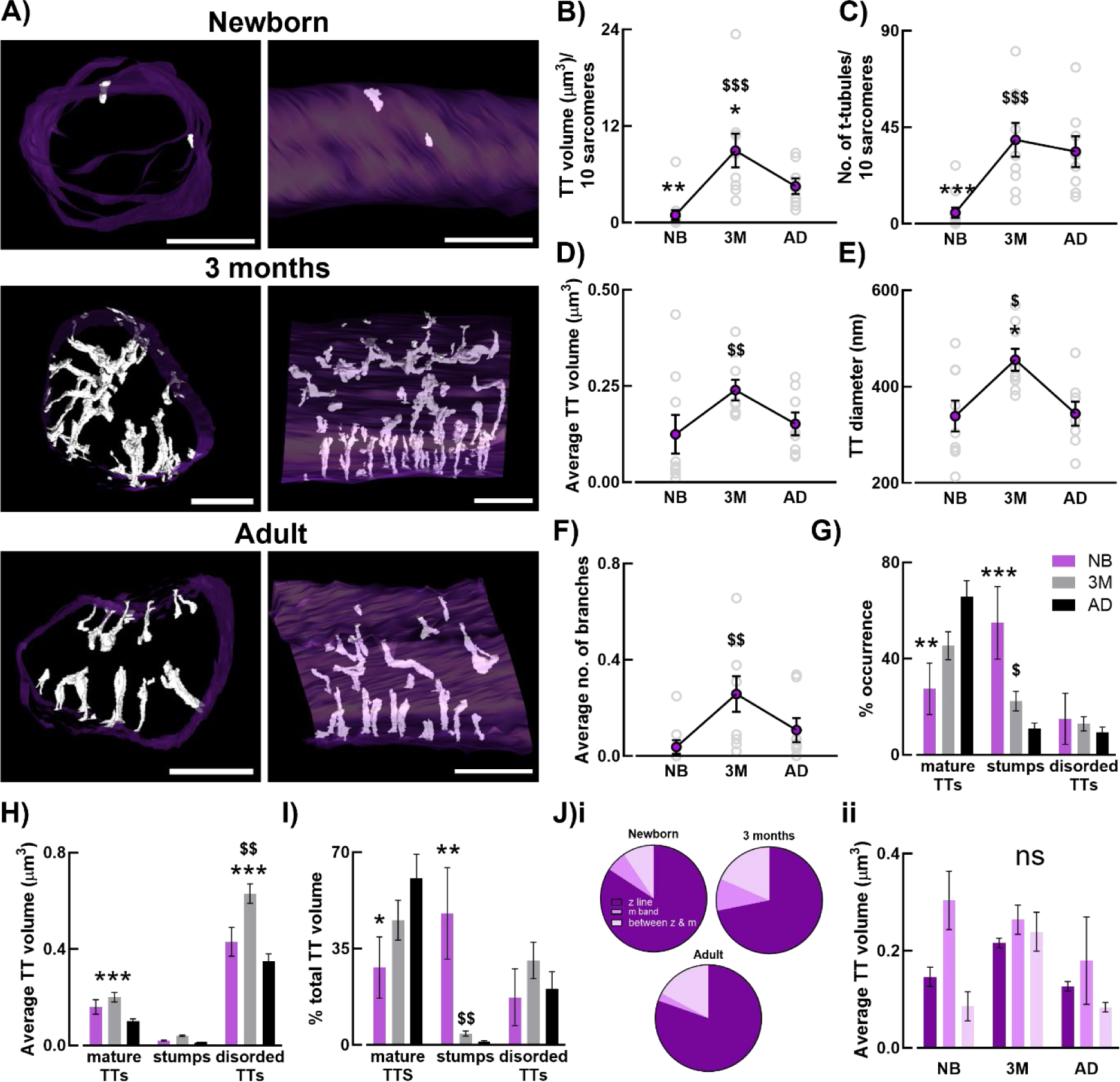
Developmental changes in t-tubule ultrastructure. Representative electron microscopy reconstructions showing the surface sarcolemma (purple) and t-tubules (white) in newborn, 3 months and adult cells (A). Mean data showing t-tubule density (B), number of t-tubules per cell (C), average t-tubule volume (D), t-tubule diameter (E), and average number of branches (F). Mean data for % occurrence (G), average volume (H) and % of total t-tubule volume for t-tubule morphologies categorised as mature (column, intermediate and club), stumps or disordered (random, oak tree, longitudinal). Summary data for t-tubule origin (J) showing occurrence from the z line, m band or between (i) and t-tubule volume depending on origin (ii). n=8-12 cells/N=3 animals per group. * *p*<0.05, ** *p*<0.01, *** *p*<0.001 *vs* adult, $ *p*<0.05, $$ *p*<0.01, $$$ *p*<0.001 *vs* previous time point, using one way ANOVA or two way ANOVA where appropriate. Scale bars = 5 µm.

**Figure 3.**
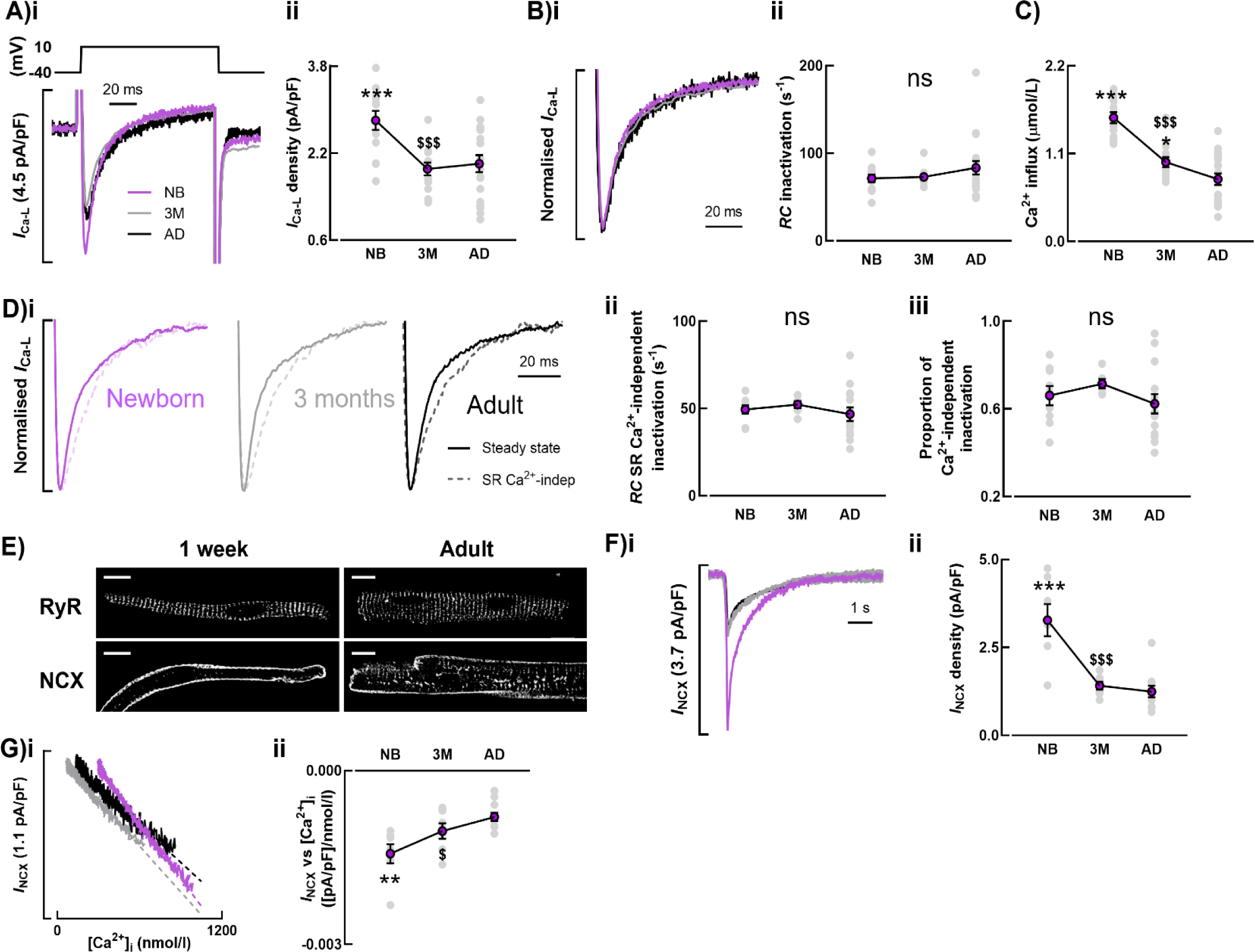
Developmental changes in *I*_Ca-L_ and *I*_NCX_. Representative traces for *I*_Ca-L_ density (Ai) and inactivation (Bi) from newborn, 3 month and adult cells. Mean data for *I*_Ca-L_ density (Aii), rate of inactivation (Bii) and Ca^2+^ influx via *I*_Ca-L_ (C). Representative *I*_Ca-L_ traces (Di) and mean data from steady state (full line) and following caffeine application (dotted line) showing the rate of Ca^2+^-independent inactivation (Dii) and proportion of Ca^2+^-independent inactivation (Diii). Representative images of RyR and NCX staining in newborn and adult cells (E). Representative traces for *I*_NCX_ density (Fi) and relationship between *I*_NCX_ and [Ca^2+^]_i_ (Gi) obtained following application of 10 mM caffeine. Mean data showing *I*_NCX_ density (Fii) and slope of the relationship between *I*_NCX_ and [Ca^2+^]_I_ (Gii). A-C, n=31-45 cells/N=13-19 animals. D, n=17-21 cells/N=6-13 animals. F&G, n=14-17 cells/N=7-11 animals. * *p*<0.05, ** *p*<0.01, *** *p*<0.001 *vs* adult, $ *p*<0.05, $$$ *p*<0.001 *vs* previous time point, using linear mixed models analysis. Scale bars = 10 µm.

Consistent with t-tubule growth from small invaginations, short stumps were most common in the newborn and decreased throughout development whereas mature t-tubule morphologies increased, consistent with t-tubule elongation (Figure 2G). This pattern is reflected in the percentage of the total t-tubule volume these structures account for, with the contribution of stumps decreasing with development but the contribution of mature t-tubules increasing (Figure 2I). The presence of stumps at all time points is consistent with continued t-tubule turnover throughout life. The number of disordered morphologies was low (Figure 2G) but since they were larger (increased volume; Figure 2H) they made significant contributions to the overall t-tubule volume (Figure 2I). Consistent with the overall t-tubule complexity and increased t-tubule volume and branching at 3 months (Figure 2A,D&F), we observed an increase in the volume of disordered morphologies at 3 months (Figure 2H).

We investigated if the complexity or increased volume of t-tubules at 3 months was due to t-tubules arising from the sarcolemma at points other than the z-line (Figure 2J). The number of t-tubule arising from a non-z-line location increased at 3 months (Figure 2Ji; *p*<0.001) and may therefore play a role in the t-tubule disorder observed. However, the point of t-tubule origin did not explain the large t-tubule volume at 3 months (Figure 2Jii).

In the ventricle, previous studies have shown that t-tubule growth is associated with increases in the Ca^2+^ transient amplitude and *I*_Ca-L_, and decreased *I*_NCX_ [15, 23, 24, 26–30]. As such, we next sought to investigate whether atrial t-tubule growth in development was associated with changes in *I*_Ca-L_ and *I*_NCX_.

### 3.4 Elevated *I*_Ca-L_ and *I*_NCX_ density at birth decrease in early life as t-tubules develop

T-tubule growth during the first 3 months of life was associated with a *decrease* in the density of *I*_Ca-L_, from 3.02±0.16 pA/pF in newborn to 1.91±0.15 pA/pF at 3 months which was then unaltered into adulthood (Figure 3Ai&ii; *p*<0.001). Decreased *I*_Ca-L_ density occurred due to increased cellular capacitance during the t-tubule development stage (Supplemental Figure 2D) which predominated over the increase in pA of *I*_Ca-L_ (Supplemental Table 1, *p*<0.001). Our data suggests an increased abundance of LTCCs at the membrane in the newborn *prior to t-tubule formation*. As t-tubules develop up to 3 months, LTCCs continue to increase however those incorporated into t-tubules likely originate from both the surface membrane and as newly formed channels. Interestingly the rate of *I*_Ca-L_ inactivation was unaltered during development (Figure 3Bi-ii, *p*=0.28) suggesting Ca^2+^ dependent inactivation was unchanged. In the newborn the increase in peak *I*_Ca-L_ and lack of change in inactivation gave rise to an increase in the integrated *I*_Ca-L_ which decreased to adulthood (Figure 3C; *p*<0.001).

We next investigated the coupling between *I*_Ca-L_ and SR Ca^2+^ release. By measuring *I*_Ca-L_ decay on the beat immediately following caffeine application, when the SR was empty, we determined the rate of decay of *I*_Ca-L_ when Ca^2+^-dependant inactivation was minimal (Figure 3Di; dotted line and 3Dii) and compared this to the rate of decay of *I*_Ca-L_ under steady state conditions (Figure 3D; solid line). The rate of inactivation on the beat immediately following caffeine application was expressed as a fraction of the steady state inactivation to estimate the proportion of *I*_Ca-L_ decay due to Ca^2+^ dependant inactivation which was unchanged in development (Figure 3Diii; *p*=0.21). Thus *I*_Ca-L_ appears similarly coupled to SR Ca^2+^ release at all developmental stages. As we have shown previously [9], RyRs adopt a z-line arrangement at birth (Figure 3E). We were unable to confirm the exact localisation of LTCCs (due to lack of a suitable antibody) but if localization is similar to that for NCX our data is consistent with both surface and t-tubule *I*_Ca-L_ existing in dyads throughout development.

Finally, in a similar way to *I*_Ca-L_, *I*_NCX_ density obtained following caffeine application (Figure 3Fi-ii), and activity for a given [Ca^2+^]_i_ (Figure 3Gi-ii), decreased to 3 months and was then unchanged to adulthood (*p*<0.001 and *p*<0.01 respectively). We next investigated if decreased *I*_Ca-L_ in development caused a decrease in the systolic Ca^2+^ transient.

### 3.5 Despite increased *I*_Ca-L_ in the newborn, Ca^2+^ transient amplitude was unaltered during development

Typical Ca^2+^ transients recorded in newborn, 3 month and adult cells are shown in Figure 4A. Despite decreased *I*_Ca-L_ and *I*_NCX_ density during development there was no difference in the amplitude of the systolic Ca^2+^ transient during development (Figure 4Bi, *p*=0.67). Diastolic Ca^2+^ decreased from newborn to 3 months while systolic Ca^2+^ was unaltered during development (Figure 4Bii-iii, *p* <0.01 and *p*=0.75 respectively). Despite decreased *I*_NCX_ during development the decay of the Ca^2+^ transient was unaltered (Figure 4Ci-ii; *p*=0.19). We next performed experiments to establish if the lack of change in Ca^2+^ transient amplitude or rate of decay were due to changes in SR Ca^2+^ content or the rate of SERCA mediated Ca^2+^ uptake.

**Figure 4.**
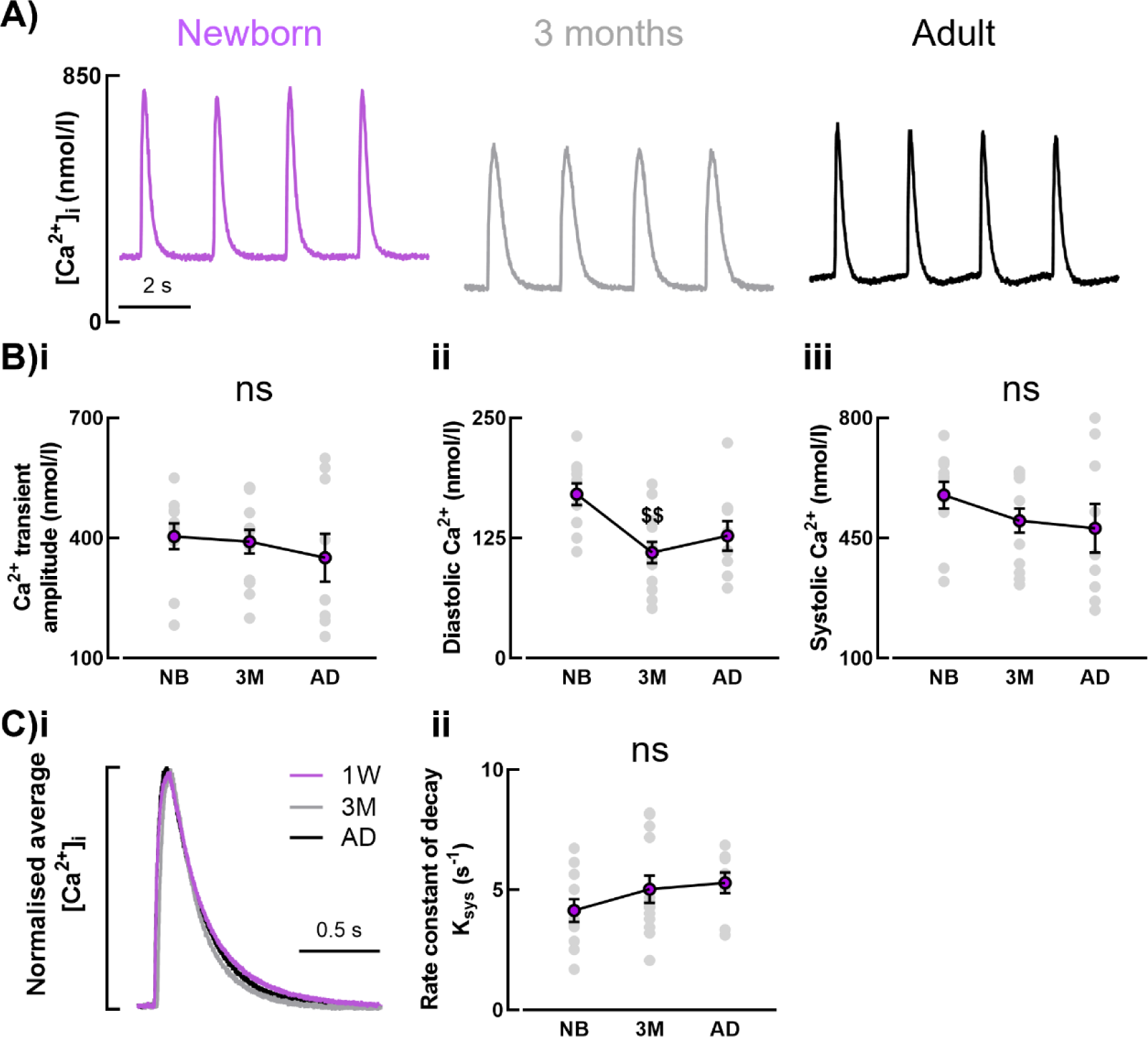
Developmental changes the systolic Ca^2+^ transient. Representative Ca^2+^ transients from newborn, 3 months and adult (A). Mean data for amplitude (Bi), diastolic (ii) and systolic Ca^2+^ (iii). Normalised average Ca^2+^ transients (i) and mean data (ii) showing rate of decay of the transient (C). n=18-35 cells/N=11-14 animals. $$ *p*<0.01 *vs* previous time point, using linear mixed models analysis.

### 3.6 Increased SR Ca^2+^ content is offset by increased cytosolic Ca^2+^ buffering

SR Ca^2+^ content was determined by application of 10 mM caffeine (Figure 5A). The amplitude of the caffeine-evoked Ca^2+^ transient was increased in the newborn which is indicative of an increase in SR Ca^2+^ content (Figure 5B; *p*<0.05). The SR Ca^2+^ content was calculated by integrating caffeine evoked *I*_NCX_ and correcting for Ca^2+^ removed by other pathways (Figure 5Aii-iii, Supplemental Figure 3). SR Ca^2+^ content was high in the newborn and decreased to 3 months and adult (Figure 5C; *p*<0.001). Therefore, a decrease in SR load cannot offset the decrease in *I*_Ca-L_ to explain the lack of change in Ca^2+^transient amplitude during development. We next examined if changes to fractional release during development could help maintain the Ca^2+^ transient. The amplitude of the systolic transient was expressed as a fraction of the amplitude of the caffeine evoked Ca^2+^ transient (Figure 5D-E). Fractional release was lower in the newborn and increased to 3 months of age (*p*<0.05; Figure 5E, *p*=0.06 overall) which would be expected to help maintain systolic Ca^2+^ despite decreases in *I*_Ca-L_ and SR Ca^2+^ content. Since decreases in *I*_Ca-L_ and SR Ca^2+^ content would both be expected to decrease fractional release, we calculated if a decrease in the ratio (SR Ca^2+^ content: *I*_Ca-L(peak)_) could explain the increase in fractional release. We found SR Ca^2+^ content decreased more than *I*_Ca-L_ during development consistent with increased fractional release (45.8 ± 5.3% *vs* 31.4 ± 5.9% respectively). We next set out to determine if, together with increased fractional release, decreased Ca^2+^ buffering could also contribute to the maintenance of the Ca^2+^ transient during development in the face of decreasing *I*_Ca-L_ and SR Ca^2+^ content.

**Figure 5.**
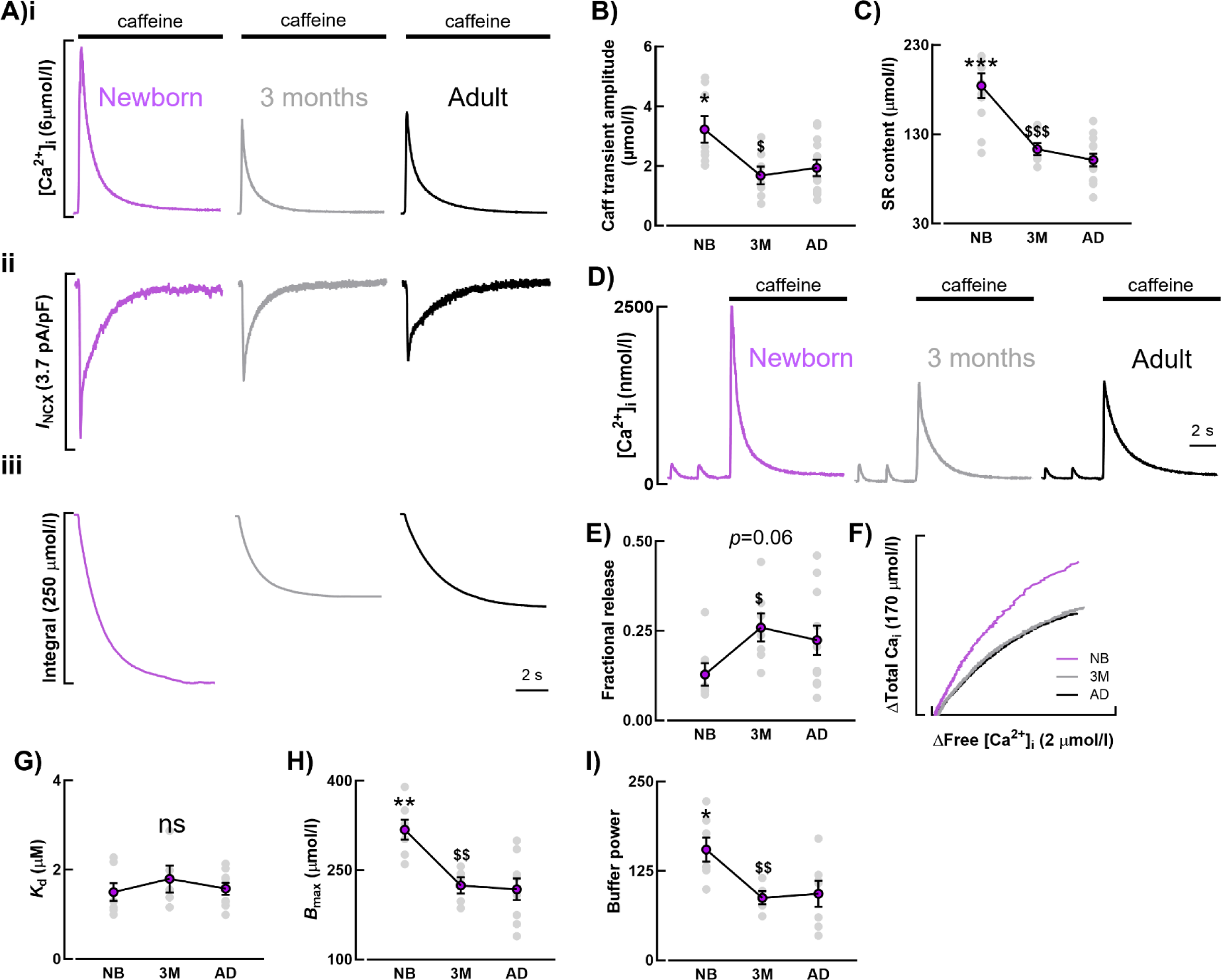
Developmental changes in SR load and Ca^2+^ buffering. Example traces following application of 10 mM caffeine (A) showing the subsequent caffeine-evoked Ca^2+^ transient (i), *I*_NCX_ (ii), and integral of *I*_NCX_ used to determine SR content (iii). Mean data for caffeine transient amplitude (B) and SR content (C). Representative Ca^2+^ transients and caffeine-evoked transients from newborn, 3 months and adult (D) reflecting the mean data for fractional release (E). Representative buffer curves from newborn, 3 months and adult (F). Mean data for buffer affinity (G), max buffer capacity (H) and buffer power (I). B, n=15-18 cells/N=7-12 animals. C, n=19-29 cells/N=7-13 animals. E, n=14-16 cells/N=7-11 animals. G&H, n=10-14 cells/N=5-9 animals. I, n=8-13/N=5-7 animals. * *p*<0.05, ** *p*<0.01, *** *p*<0.001 *vs* adult, $ *p*<0.05, $$ *p*<0.01, $$$ *p*<0.001 *vs* previous time point, using linear mixed models analysis.

Ca^2+^ buffer curves were constructed by plotting the change in total Ca^2+^, calculated from the *I*_NCX_ integral, as a function of the change in free Ca^2+^, measured from the caffeine evoked Ca^2+^ transient and were fit with a hyperbolic function (equation 1; Figure 5F). In the newborn the buffer curve was shifted to the left, indicative of an increase in Ca^2+^ buffering which decreased to adult levels by 3 months (Figure 5F). The affinity of buffers for Ca^2+^ (*K*_d_) was unchanged during development (Figure 5G; *p*=0.41) but maximum buffering capacity (*B*_max_) was high in the newborn and decreased at 3 months and adult (Figure 5H; *p*<0.01). Buffer power, calculated using *K*_d_ and *B*_max_ in equation 2 as previously published [33], was also high in the newborn and decreased at 3 months and adulthood (Figure 5I, *p*<0.05). Decreased Ca^2+^ buffering requires a smaller change in total Ca^2+^ to produce the same change in free Ca^2+^ and could therefore play a role in the lack of change in Ca^2+^ transient amplitude despite decreases in *I*_Ca-L_ and SR Ca^2+^ content during development. To remove effects of Ca^2+^ buffering we next examined the properties of the total Ca^2+^ transient to determine the contribution of Ca^2+^ buffering in setting the properties of the systolic Ca^2+^ transient.

### 3.7 Decreased Ca^2+^ buffering maintains the amplitude of the systolic Ca^2+^ transient during development

The mean total Ca^2+^ transient at each developmental time point is shown in Figure 6A. It is evident that the amplitude of the total Ca^2+^ transient decreases from newborn to 3 months and adult consistent with decreases in *I*_Ca-L_ and SR Ca^2+^ content suggesting it is the decrease in Ca^2+^ buffering which maintains the free Ca^2+^ transient during development. Similarly to the rate of decay of free Ca^2+^, the rate of decay of total Ca^2+^ was unaltered during development (Figure 6Aiii). However, the abundance of SERCA and the ratio between SERCA and phospholamban (PLN) decreased during development suggesting other factors play a role.

**Figure 6.**
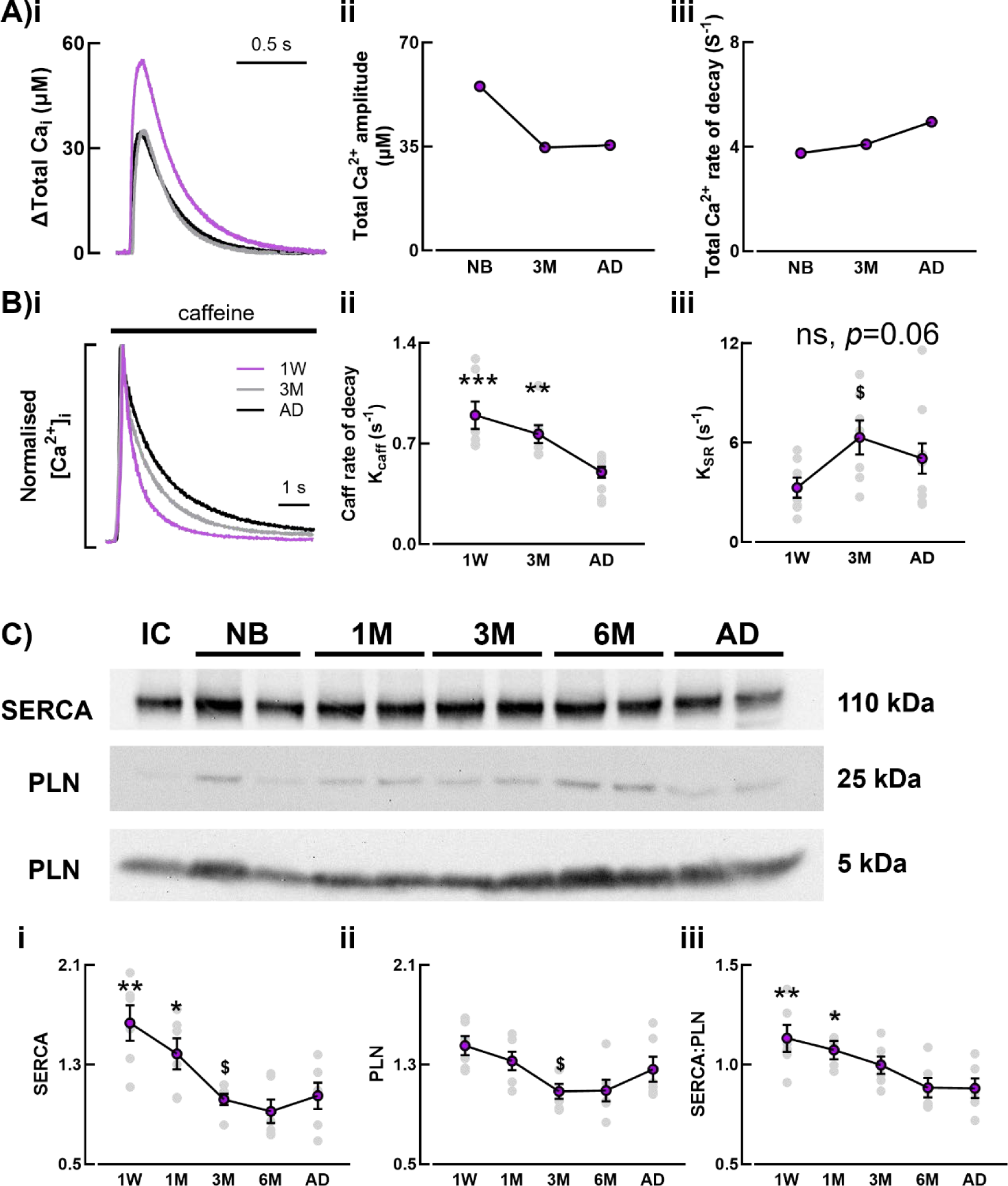
Decreased Ca buffering maintains the amplitude of the Ca^2+^ transient during development. Ai) Mean total Ca^2+^ transients were generated from the mean free Ca^2+^ transient (n=18-35 cells/N=11-14 animals) using the average buffering parameters at each developmental time point and equation 1 (n=10-14 cells/N=5-9 animals). Ai-ii show the amplitude and rate of decay of the mean total Ca^2+^ transients in (Ai). The abundance of SERCA and PLN at each developmental time point was calculated by western blot. Typical membranes probed for SERCA and PLN (monomer and pentamer) are shown in Panel Bi-iii show how SERCA, PLN and the SERCA:PLN ratio change with development. B, n=13-16 cells/7-10 animals. C, N=6 animals, minimum 3 repeats. * *p*<0.05, ** *p*<0.01, *** *p*<0.001 *vs* adult, $ *p*<0.05 *vs* previous time point, using linear mixed models analysis.

## 4. Discussion

In this study we show that the time course for postnatal t-tubule development and Ca^2+^ handling changes in the atria is in stark contrast to that in the ventricle. We found that when t-tubules are sparse in early life, increased *I*_Ca-L_ and SR load help maintain the Ca^2+^ transient by overcoming increased intracellular Ca^2+^ buffering. As the atrium matures postnatally, t-tubules adopt a specialist developmental state characterized by increased t-tubule density and complexity. Through the transition to adulthood, greater triggered Ca^2+^ release mediated by increased t-tubule density and decreased Ca^2+^ buffering permits maintenance of Ca^2+^ transient amplitude despite reduced *I*_Ca-L_ and SR load.

### 4.1 Cardiac growth is accelerated in the early postnatal period and the adult phenotype is attained at 3 months of age

Accelerated cardiac growth was observed in the early postnatal period, indicated by HW:BW. This has been previously reported in multiple species[50, 51] and is facilitated by hyperplastic followed by hypertrophic growth. The trigger for this has not been fully defined but is attributed to increased arterial pressure[52] and hormone changes[53, 54] amongst others. Little is currently known about hypertrophic growth in the atria, but the switch to terminal differentiation in the ventricle appears to be species-dependent, occurring *in utero* in human and postpartum in rodents[20, 55]. As some binucleated atrial cells were present in the neonate this indicates this switch begins in utero as in the ventricle; though as 80% of sheep ventricular myocytes are binucleated at birth[48] this suggests chamber differences in gross maturation. Indeed, as increases in binucleation, cell size, capacitance and volume continued to be observed during development it is possible that there is a slower rate of growth in the atria *vs* ventricle.

Interestingly, the plateauing of changes in HW:BW ratio at 3 months coincides with maximal t-tubule density being attained. Since the function of t-tubules is to facilitate triggered Ca^2+^ release throughout the cell, it is perhaps unsurprising that t-tubules are in situ for the period of hypertrophic growth.

### 4.2 Atrial t-tubules enter a specialist state during development

In contrast to small mammalian species [15–18, 21], we and others have previously shown the presence of t-tubules in foetal and newborn sheep[9, 19] and human foetal [20] ventricles. The presence of t-tubules at birth may be dependent on gestation period and developmental maturity with the time course for t-tubule formation shifted but the process and subsequent maturation of EC coupling appearing similar between species. This may be correlated with the shift from hyperplastic to hypertrophic growth previously discussed and is suggested to be associated with changes in cortisol and thyroid hormone that occur *in utero* in large mammals[53, 54]. This is supported by glucocorticoid and thyroid hormones promoting t-tubule development in human-induced pluripotent stem cell-derived cardiac myocytes[56].

Despite differences in the time course for development, previous studies in the ventricle appear to show continual t-tubule formation into adulthood, with positive correlation between t-tubule density and cell size[15, 16, 19]. It is unclear as to whether this is true or if there is also an overshoot in t-ventricular tubule formation prior to adult, as in the atria, due to the limited time points studied. Whilst this is unknown, our study presents a novel finding that t-tubule formation is not directly associated with changes in cell width despite adult atrial t-tubule density being positively correlated to cell width across species[3]. At present, the cause of this overshoot is unclear. Though we found no clear association with proteins known to be involved in t-tubule formation in the ventricle[2, 15, 19], our observed decrease in BIN1 expression is similar to that shown after postnatal day 15 in mice. Given the species differences in the time course for t-tubule development, this likely reiterates a role for BIN1 in early t-tubule formation with expression subsequently maintained in line with continued t-tubule turnover as evidenced by the presence of t-tubule stumps at all time points.

Whilst this warrants further investigation, it is interesting that the increased t-tubule density we observe at 3 months is due to differences in t-tubule structure not number. Compared to the ventricle[43], atrial t-tubule structures are more diverse and complex, with a number of non-uniform shapes observed. The reasons behind this are not understood but as atrial t-size and complexity are increased at 3 months, it is possible that the initial demand for t-tubule growth results in disordered structures that are trimmed and optimised for function in adult. Indeed, as CICR is not enhanced at 3 months despite increased t-tubule density, the t-tubule network appears relatively inefficient *vs* adult.

### 4.3 *I*_Ca-L_ and SR load maintain Ca^2+^ transient amplitude in newborn to overcome increased Ca^2+^ buffering and reduced coupling with the SR

In contrast to the ventricle where *I*_Ca-L_ increases with t-tubule maturation[15, 24, 26], *I*_Ca-L_ in the atria was maximal in the neonate when t-tubules were sparse. Whilst *I*_Ca-L_ is concentrated on t-tubules in the adult ventricle[57], there is even distribution on the surface and t-tubules in the atria[58] which may contribute to the disparity observed between chambers during development. Despite this, the increased newborn *I*_Ca-L_ should act as a greater trigger for SR release[36], yet alongside increased SR load we found no difference in systolic Ca^2+^ transient amplitude. We have shown this previously [9]and it has also been shown in the human atria[59], in contrast to studies in the ventricle where the amplitude of the transient increases postnatally[15, 17], despite reports of increased SR load in early life[60]. This preservation of atrial Ca^2+^ transient amplitude indicates atrial function is important throughout development, and may be compensatory for reduced ventricular relaxation that has been reported in early life[61–63]. As the atria contribute 25% to ventricular filling[64], it is possible that they may play a key role in ensuring cardiac output is maintained in the neonate and sustained into adulthood.

As indicated in our study, functional SR stores exist in the neonate human atria but that the efficiency of release increases with age, likely as t-tubules develop[59]. We have previously shown RyR clusters are aligned at the z-line from birth and are enlarged compared to adult [9]. Given fewer t-tubules in early life leave SR in the cell centre uncoupled, reduced triggered central Ca^2+^ release may not only contribute to the unaltered Ca^2+^ transient we observe but also to the increased SR load, along with increased Ca^2+^ influx[65] and SERCA:PLN expression. In the neonate rabbit ventricle reverse-mode operation of NCX compensates for both reduced *I*_Ca-L_ and SR coupling[27–30]. As we saw increased *I*_Ca-L_ in the neonate atria we suggest that the increased *I*_NCX_ we also observed is reflective of increased operation in forward-mode to ensure efflux equals influx.

Alongside reduced coupling with the SR, we attribute unaltered Ca^2+^ transient amplitude to increased buffering. As previously discussed in [33, 41], 99% of cytosolic Ca^2+^ is bound to intracellular buffers. With buffering increased in the neonate, a greater amount of total Ca^2+^ is required to generate a rise in free Ca^2+^ similar to adult. We found no difference in buffer *K*_d_ in the study but increased *B*_max_ indicating the endogenous buffers are consistent throughout development, but that expression is increased in the newborn. This could be attributed to the greater SERCA expression we observed in early life or potential changes in troponin that have not been examined here[66]. Whist increased buffering alone could account for the lack of change in Ca^2+^ transient amplitude, it may also exacerbate impaired CICR due to reduced coupling in early life as it limits propagation that is required to facilitate central Ca^2+^ release in the absence of a developed t-tubule network[67].

Comparisons are often made between the neonate and the failing heart to understand how myocytes can function with reduced t-tubule density in health but not disease. We and others show efficient *I*_Ca-L_ coupling despite immature t-tubules in early life, with improved coupling with SR throughout the cell as t-tubules develop[15, 18]. In HF, t-tubule loss in the ventricle results in a return to impaired central Ca^2+^ release, however EC coupling is also diminished at structurally coupled sites that are functional in the neonate[18]. Though no direct comparisons have been made between the neonate and HF atria in this study, it is likely that t-tubule remodelling is compounded by other factors resulting in impaired CICR in failing atrial cells. This may include the observed reduced *I*_Ca-L_ triggering less Ca^2+^ release even from SR close to the surface[13, 36], and a possible role for remodelling of RyRs[18, 68].

### 4.5 Conclusions

Our data shows that when t-tubules are sparse in the newborn atria, myocytes are specially adapted to maintain Ca^2+^ transient amplitude with increased *I*_Ca-L_ and SR load compensating for increased buffering. Despite reductions in *I*_Ca-L_ and SR load with age, Ca^2+^ transient amplitude is maintained due to increased t-tubule density, triggered central Ca^2+^ release and reduced Ca^2+^ buffering. Given the preservation of the Ca^2+^ transient ensures maintained atrial function, the neonate atria can provide insights into adaptations in Ca^2+^ cycling to negate reduced t-tubule density that occurs in HF.

## Author Contributions

All authors contributed to the delivery of experiments/data analysis. KMD, AWT and DAE obtained funding and were responsible for the experimental concept. CERS and KMD wrote the manuscript, and all authors contributed to manuscript editing.

## Acknowledgments

The authors thank the staff in the FBMH EM Core Facility for their assistance and the Wellcome Trust for equipment grant support to the EM Facility.

---

**Supplemental Table 1.** Additional cell parameters. Mean additional cell data. For length and volume n=31-28 cells/N=3-5 animals, for raw peak *I*_Ca-L_, n=15-45 cells/N=7-19 animals. Statistical significance determined using mixed model analysis.

| Time point | Cell length (μm) | Cell volume (pL) | Raw peak $I_{Ca-L}$ (pA) |
| --- | --- | --- | --- |
| Newborn | 91.7±3.2 | 4.6±0.2 | 77.8±7.4 |
| 1 month | 124.3±11.8 | 9.4±2.0 | 111.1±10.4 |
| 3 months | 132.2±1.4 | 13.0±0.7 | 135.4±10.63 |
| Adult | 133.9±9.1 | 16.0±2.3 | 136.6±11.3 |
| | *** $p<0.001$ | *** $p<0.001$ | *** $p<0.001$ |

**Supplemental Figure 1.**
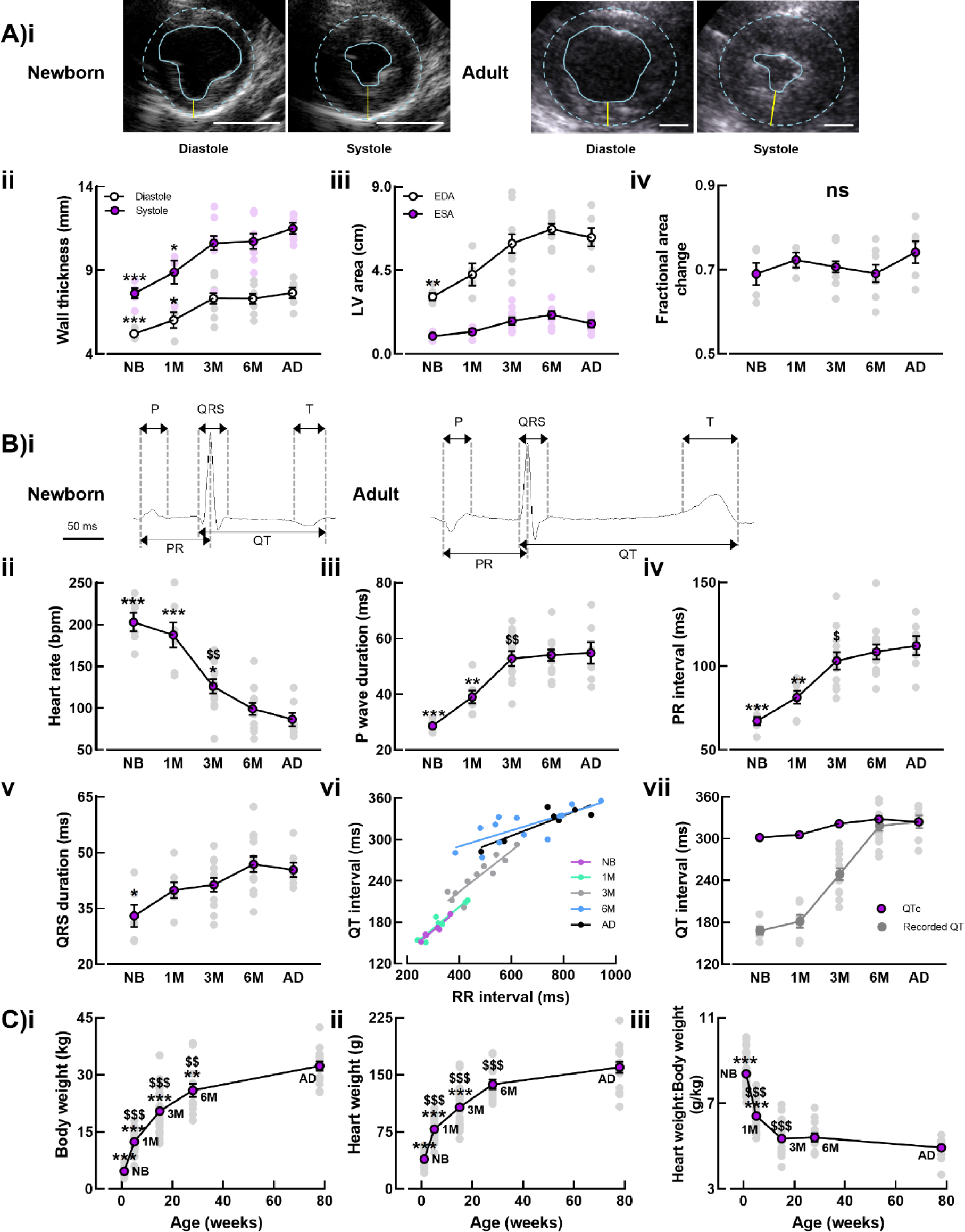
In vivo characterisation of sheep postnatal development. Transthoracic echocardiography was used to measure developmental changes in ventricular size and contractile function. (Ai) Representative short axis views during diastole and systole of the newborn and adult heart. Posterior wall thickness (Aii, yellow line Ai) and left ventricular area (end diastolic area; EDA and end systolic area; ESA; Aiii). Fractional area change (Aiv) calculated as the (ESA – EDA)/EDA. Typical electrocardiograms for the newborn and adult (Bi) that were used to calculate heart rate (Bii) and mean ECG parameters (Biii-vii) measured as shown in Bi. Recorded QT interval (grey symbols) and the corrected QT interval (QTc) where QT was corrected for heart rate using the regression lines displayed in Bvi (Bvii). Mean body weight, heart weight and the heart weight:body weight ratio calculated over the time course of neonatal development (Ci-iii). A, N=4-10 animals. B=6-13 animals. C, N=15-40 animals. * *p*<0.05, ** *p*<0.01, *** *p*<0.001 *vs* adult, $ *p*<0.05, $$ *p*<0.01, $$$ *p*<0.001 *vs* previous time point, using one way ANOVA. Scale bars = 2 cm.

**Supplemental Figure 2.**
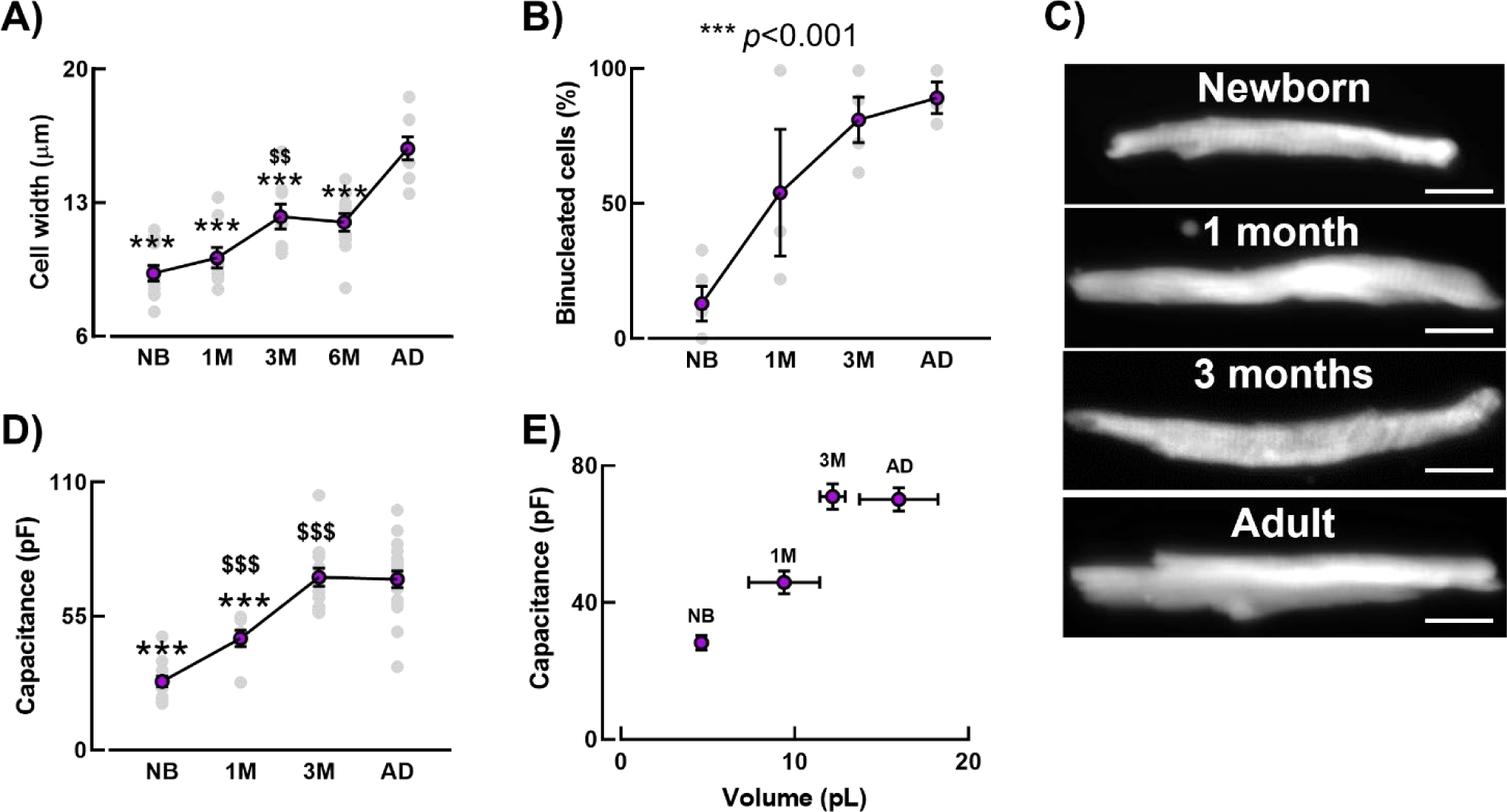
Developmental changes in atrial myocyte size and structure. Mean data for cell width obtained from confocal z-stacks of cells stained with ANEPPS (A). Cells at each developmental stage were loaded with calcein and imaged confocally through the z-plane (C). The number of nuclei were also recorded from the calcein images and expressed as the percentage of binucleated cells (B). A 10 mV hyperpolarising step was applied to patch clamped cells to record capacitance (D). Capacitance was plotted against cell volume, calculated from confocal z-stacks where three dimensional reconstructions, to calculate surface area to volume ratios and the relationship shown in E. A, n=41-61 cells/N=8-11 animals For capacitance, n=15-45 cells/N=7-19 animals, for volume, n=29-38 cells/N=3-5 animals. * *p*<0.05, ** *p*<0.01, *** *p*<0.001 *vs* adult, $ *p*<0.05, $$ *p*<0.01, $$$ *p*<0.001 *vs* previous time point, using linear mixed models analysis. Scale bars = 10 µm.

**Supplemental Figure 3.**
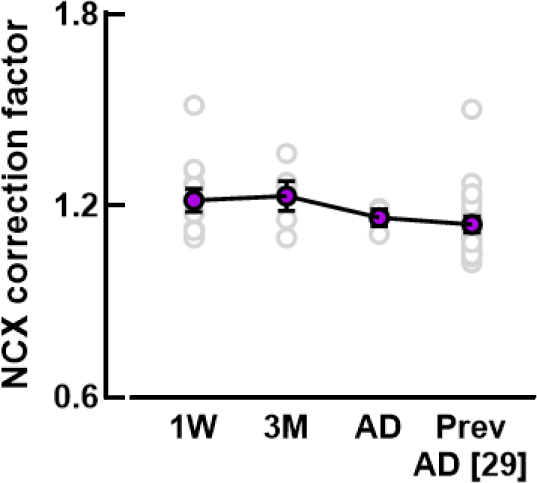
NCX correction factor is unchanged in development. The contribution of pathways other than *I*_NCX_ to Ca^2+^ removal during caffeine application was calculated by calculating the rate of decay of the caffeine evoked Ca^2+^ transient in the presence of 10mM Nickel (to inhibit NCX) and is displayed with our previously published value. n=3-20 cells/N=2-8 animals.

